# Engineering the soil bacterium *Pseudomonas synxantha* 2-79 into a ratiometric bioreporter for phosphorus limitation

**DOI:** 10.1101/2023.10.20.563366

**Authors:** Elin M. Larsson, Richard M. Murray, Dianne K. Newman

## Abstract

Microbial bioreporters hold promise for addressing challenges in medical and environmental applications. However, the difficulty of ensuring their stable persistence and function within the target environment remains a challenge. One strategy is to integrate information about the host strain and target environment into the design-build-test cycle of the bioreporter itself. Here, we present a case study for such an environmentally-motivated design process by engineering the wheat commensal bacterium *Pseudomonas synxantha* 2-79 into a ratiometric bioreporter for phosphorus limitation. Comparative analysis showed that an exogenous P-responsive promoter outperformed its native counterparts. This reporter can selectively sense and report phosphorus limitation at plant-relevant concentrations of 25-100 *µ*M without cross-activation from carbon or nitrogen limitation or high cell densities. Its performance is robust over a field-relevant pH range (5.8-8), and it responds only to inorganic phosphorus, even in the presence of common soil organic P. Finally, we used fluorescein-calibrated flow cytometry to assess whether the reporter’s performance in shaken liquid culture predict its performance in soil, finding that although the reporter is still functional at the bulk level, its variability in performance increases when grown in a soil slurry as compared to planktonic culture, with a fraction of the population not expressing the reporter proteins. Together, our environmentally-aware design process provides an example of how laboratory bioengineering efforts can generate microbes with greater promise to function reliably in their applied contexts.

## Introduction

In the past 30 years, whole-cell microbial bioreporters and biosensors have been developed for applications in environmental sustainability and medicine. In the simplest case, they are made up of a promoter that gets activated by a target signal that then drives the expression of genes that result in a measurable output such as luminescence or fluorescence. The simplicity and versatility of this structure has allowed the development of biosensors for heavy metals [1], pathogens [2] and plant nutrients [3].

One advantage of microbial biosensors is that they can detect the target analyte concentrations at microscopic scales in an environment of interest. This makes them promising candidates for future monitoring technologies in medicine and agriculture. An application where they could be useful is in precision agriculture, a farming practice that collects spatial and temporal information about different parameters, such as moisture content and nutrient levels, and uses it to make decisions about where action needs to be taken [4]. In the case of fertilizer application, for this to be effective, the detection of nutrients must target the bioavailable portion that can be utilized by crops. For this reason, detection methods that measure the total nutrient content are not as informative. If high resolution, accurate measurements can be made of bioavailable nutrient concentrations, that information can be used to apply an appropriate amount of fertilizer, reducing the over-application. A good target nutrient for this application is phosphorus, a non-renewable resource commonly added to agricultural fields. When applied to soils it can get bound by minerals, which makes it unavailable for plant uptake [5]. It is also common for the phosphorus to be flushed away with surface run-off, ending up in water bodies where it causes eutrophication.

A prerequisite for engineering a microbial biosensor for agriculture is the ability of the chassis to stably colonize the soil environment and persist throughout the growing season. For example, *E. coli* K-12 gets outcompeted rapidly, about two weeks after being introduced to soil [6]. Pseudomonads, a class of bacteria that are ubiquitous in soil, have been common target chassis for engineering soil biosensors. Not only are they well-adjusted to the soil environment, they are in many cases known to promote plant-growth and protect against plant pathogens [7, 8]. Although it is not certain that engineered isolates can persist long-term in soil, it has been reporter previously that can persist and retain their engineered function for months in soil [9].

The first engineered bioreporter for phosphorus limitation in a pseudomonad was developed by de Weger et al. [10]: they inserted *lacZ* in random places in the *Pseudomonas putida* WCS358 genome and found colonies that responded to phosphorus limitation with *β*-galactosidase activity. A similar approach was taken by Kragelund et al. [11] who instead integrated the *lux* operon onto the *Pseudomonas fluorescens* DF57 genome. The first use of an exogenous P limitation promoter in a pseudomonad was done by Dollard et al. [12], who used the *E. coli* P_*phoA*_ promoter in *Pseudomonas fluorescens* DF57 and showed successful expression of the *lux* operon during P limitation. Native promoters have evolved in the context of maximizing host fitness [13, 14]. Sometimes non-native genetic elements can outperform native ones for engineering applications that instead prioritize sensor performance, for example by maximizing expression levels or by reducing the risk of metabolic cross-talk [15, 16]. However, to our knowledge there has not been a direct comparison between the performance of the *E. coli* P_*phoA*_ promoter and native *Pseudomonas* promoters for use in P limitation reporters. In addition, none of the described P reporters implement a ratiometric output readout. Ratiometric reporters are advantageous because they control for cellular metabolic activity and permit normalization of the bulk output signal.

In this study, we develop a ratiometric reporter for P limitation in the wheat isolate *Pseudomonas synxantha* 2-79. We selected this strain as our chassis because it is known for its ecological importance in the biocontrol of wheat in the Pacific North-west Columbia Plateau, and phosphorus content has been mapped in this region using traditional methods [7].

We characterize its performance under environmentally-relevant conditions, including pH and phosphate source, cross-talk with other nutrient limitations, and performance in a soil context. Our work builds upon previous efforts by (1) directly comparing the performance of the exogenous P_*phoA*_ promoter to native promoters, showing that the P_*phoA*_ promoter outperforms native promoters and (2) creating a ratiometric sensor.

## Results and Discussion

### Selection and characterization of a promoter induced by phosphorous limitation

We first set out to determine whether the exogenous *E. coli* P_*phoA*_ promoter performs better or worse than native *P. synxantha* promoters as a bioreporter for phosphorous (P) limitation. Specifically, we compared P_*phoA*_ to five native *P. synxantha* promoters. Three were chosen by performing RNA-sequencing and selecting the three most upregulated genes (Supplemental information A: Table S1 and Material and Methods). Additionally, two promoters that have annotated binding regions in *Pseudomonas fluorescens* Pf1-0 [17] were chosen as promoter candidates for the reporter. We used the intergenic region upstream of the five chosen genes to construct all promoter fusions (Supplemental information: Table S1).

These promoters all respond to the PhoB-PhoR two-component system to sense the limitation of phosphate in their surroundings (Figure 1A, left). After phosphate has been imported to the periplasm through porins, it passes through the phosphate transporter PstS into the cytosol [19]. When the transporter is saturated, the adjacent histidine kinase PhoR remains inactive, however, when the levels of phosphate reach a critically low level (4 *µ*M in *E. coli*) [20], PhoR is activated and phosphorylates the transcriptional regulator PhoB. The activation of PhoB leads to its binding to DNA regions called PHO-boxes that are found upstream of genes that are part of the Pho-regulon, involved in conservation and scavenging of phosphorus [19] .

**Figure 1:**
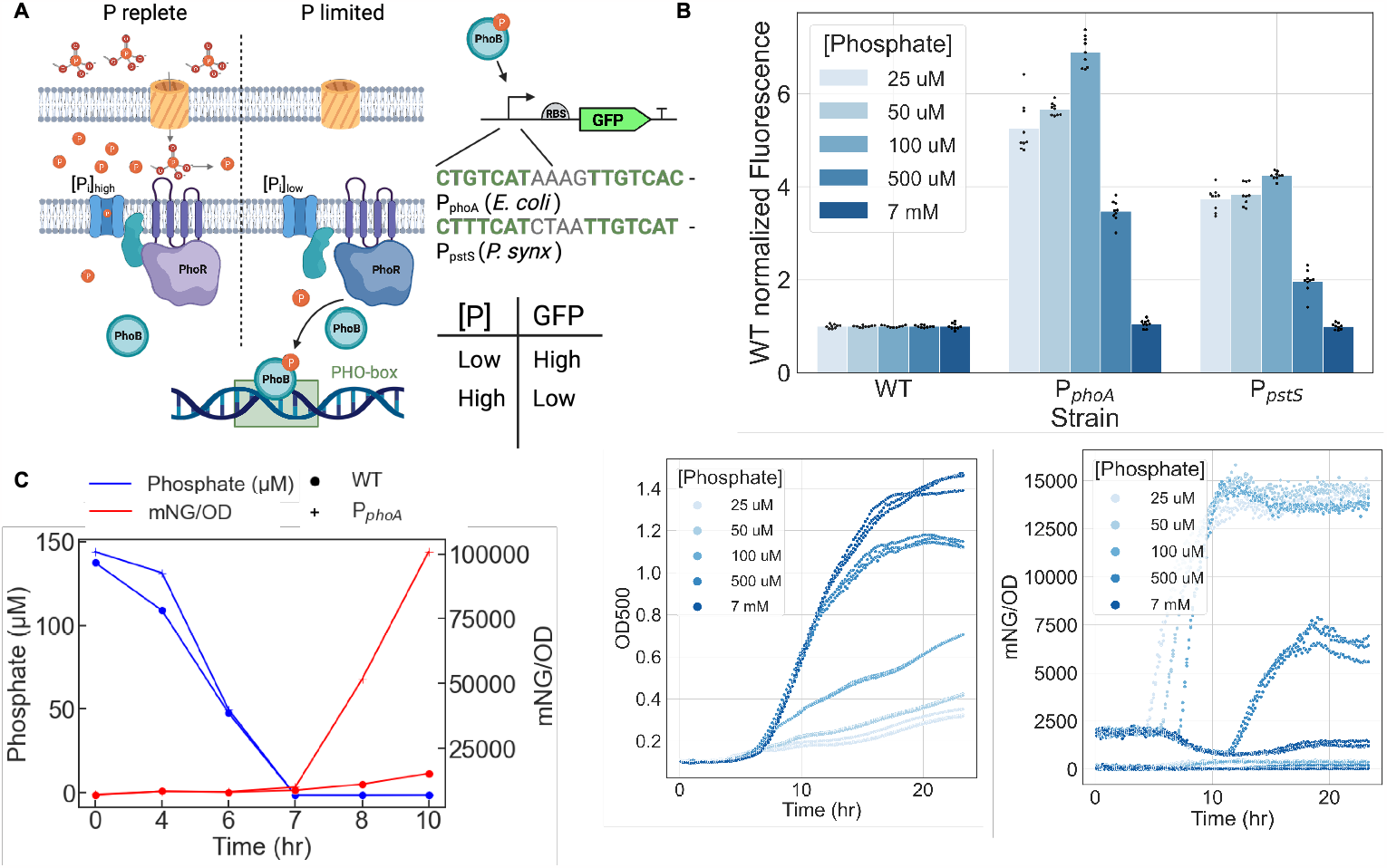
(A) Left: Schematic of *E. coli* phosphorus sensing. Right: Construct diagram for the two promoter fusions. The PhoB binding region is annotated in bold. The *E. coli* annotation is from [18]. The annotation for the *P. synxantha* promoter are from *Pseudomonas fluorescens* Pf0-1 in [17]. (B) Upper: Promoter response to different initial phosphate concentrations. The data is normalized by the WT fluorescence at the final timepoint of the experiment (24 hours). Bars show the average fluorescence value, dots show raw data for three biological replicates. Lower: Representative plots for growth and fluorescence/OD for P_*phoA*_ at different initial phosphate concentrations. The lag-time for the response is longer the more phosphate is provided at the start. (C) One out of three biological replicates showing mNG/OD (red) and phosphate concentrations (blue) over 10 hours of growth (the two other biological replicates are found in Supplemental information: Figure S3). Phosphate is depleted in the culture over time. Once a critically low concentration (below 50 *µ*M) is reached, the GFP signal is turned on at 7-8 hours of growth for the reporter cells (squares) whereas WT (circles) fluorescence increases only by a small amount.

We developed reporter constructs for the selected promoters by cloning them upstream of the green fluorescent protein mNeonGreen (mNG) [21], driven by a strong synthetic ribosome binding sequence [22] (Figure 1A, right). Each reporter variant was integrated in single copy on the *P. synxantha* genome using transposon-based integration at the Tn7 site [23]. We then grew these constructs strains under P limited (25-500 *µ*M) and P replete (7 mM) conditions and measured their optical density (OD) and fluorescence. Because *P. synxantha* makes phenazines [24] and pyoverdine [25] that can interfere with the mNG signal, we also grew wild-type (WT) cells that did not contain the reporter construct to normalize the output of the reporter cells by the native cell background.

We observed that of the five native promoters tested, P_*pstS*_ had the strongest signal (Supplemental information: Figure S1). Three of the promoter fusions had no significant signal compared to WT. The lack of signal from the P_*phoX*_ and P_*phoD*_ promoters was surprising, as reporters had previously been constructed in *Pseudomonas fluorescens* with annotated regions of the PHO-box [17]. One potential explanation could be secondary structures forming between the promoter and RBS region, hindering the ribosome from binding. An alternative explanation could be that other growth conditions than the ones tested are necessary for activation in *P. synxantha* 2-79.

After identifying P_*pstS*_ as the strongest native promoter, we compared its response to the non-native *E. coli* P_*phoA*_ promoter at a range of initial P concentrations. We observed that the promoter P_*phoA*_ had the strongest signal with a fluorescence that was 5.3-to 6.9-fold higher than the WT strain (Figure 1B). The native P_*pstS*_ promoter was 3.7-to 4.2-fold more fluorescent than WT. Both promoters showed similarly minimal levels of leak, with fluorescence signals being comparable to WT levels at replete (7 mM) P concentrations.

For the promoters that responded to P limitation, we observed that increasing P concentrations corresponded to delayed activation of the mNG signal (Figure 1B, bottom, Supplemental information: Figure S2). We hypothesized this delay arises because the cells gradually deplete the phosphorous in their medium and only turn on the mNG signal when the P concentration crosses below a critical threshold. We tested this hypothesis by simultaneously measuring the cellular fluorescence and phosphate concentration from a growing culture of the the P_*phoA*_ reporter strain (Figure 1C, Supplemental information: Figure S3). As expected, the critical threshold for mNG signal activation was below 50 *µ*M, which is consistent with the range of P concentrations that are limiting for wheat (20-200 *µ*M) [26].

Taken together, these results demonstrate that the exogenous *E. coli* promoter P_*phoA*_ has a stronger response to P limitation at physiologically relevant concentrations than two native P-responsive promoters in *P. synxantha* 2-79. We therefore chose P_*phoA*_ as the basis for the continued characterization and development of our bioreporter.

### The *E. coli* P_*phoA*_ promoter does not exhibit cross-talk with carbon and nitrogen starvation conditions

Minimizing cross-talk, the activation of the response by unintended signals, is one of the central challenges in engineering bioreporters. There are two major categories of cross-talk: it could arise from the activation of the promoter by other cellular elements responding to the target signal in unpredictable ways (genetic cross-talk), or it could arise from other signals directly activating the promoter (Figure 2A). Because we chose to use an exogenous promoter as the basis for our bioreporter, the possibility of genetic cross-talk was particularly important to address.

**Figure 2:**
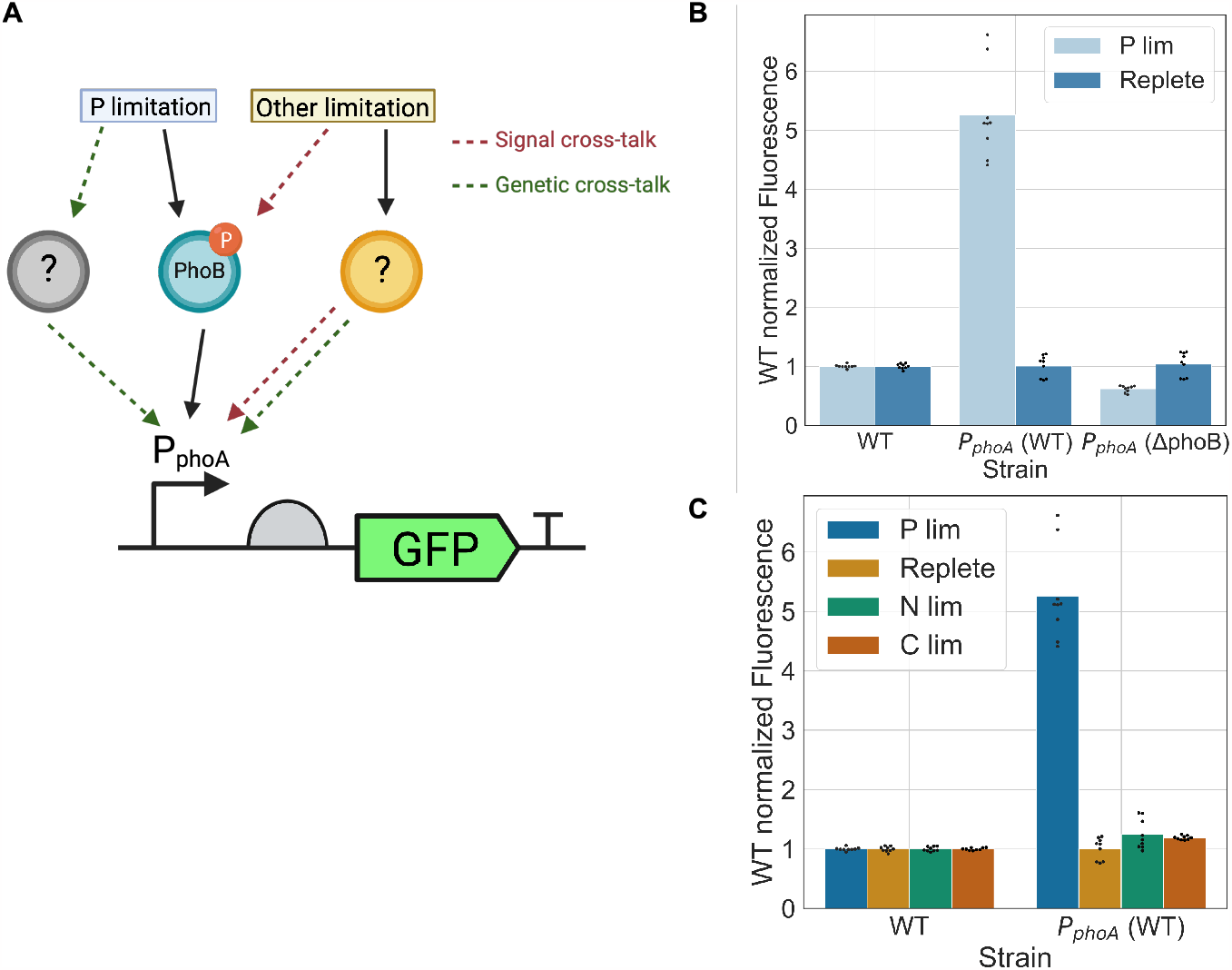
(A) Schematic showing potential genetic and signal cross-talk in the system. (B) The reporter fusion was integrated in the ΔphoB background and grown in P limited (50 *µ*M) and P replete (7 mM) media. The fluorescence output is compared to the reporter in the WT background and normalized to WT. (C) P reporter grown in medium limited for carbon, nitrogen or phosphorus compared to growth in nutrient replete medium. WT normalized fluorescence signal is plotted as an endpoint value at 24 hours of growth. Bars show the average fluorescence value, dots show raw data for three biological replicates.

To test whether the activation of P_*phoA*_ during P limitation arises exclusively through the PhoB-PhoR pathway, we integrated the P_*phoA*_ reporter into a Δ*phoB* deletion strain and measured its response to P limitation (Figure 2B). We observed that the reporter in the Δ*phoB* strain does not activate mNG expression during P limitation, indicating that PhoB is indeed necessary for P_*phoA*_ activation in *P. synxantha* 2-79.

We next sought to determine the extent of signal cross-talk in our system by assessing the reporter’s response to other types of nutrient limitation. Metabolic pathways for different nutrients are sometimes regulated by overlapping pathways in bacteria [19, 27, 28], as this can help the cell adapt to various environmental nutrient conditions [29].

We measured the response of the P_*phoA*_ reporter to limiting concentrations of phosphorous (P), carbon (C), and nitrogen (N), as these are all nutrients that can limit bacterial growth in soils [30]. The reporter had a minimal response to C and N limitation (approximately 1.2-fold average increase in fluorescence over WT) compared to P limitation (approximately 5.3-fold average increase over WT) (Figure 2C).

Together, these experiments confirm that the P_*phoA*_ promoter exhibits minimal genetic cross-talk and signal cross-talk to other relevant limiting nutrients.

### pH has variable effects on reporter performance

Another challenge associated with engineering microbial bioreporters is that environmental parameters like pH often differ from those of the standard laboratory conditions typically used to prototype the constructs. Furthermore, in natural environments like soils, the values of these parameters can vary over time and across sites [31]. The pH of the Cook Agronomy Farm, from where the *P. synxantha* strain in this study was isolated, has historically been reported to range between acidic and alkaline depending on the sampling location [32, 33, 34]. Ideally, a reporter should function robustly across a range of different pH conditions as well as changes in pH caused by biological processes.

To assess the robustness of our reporter to different pH conditions, we measured its response to P limitation under acidic (pH=5.8), neutral (pH=7), and alkaline (pH=8) conditions. We additionally measured the performance of the P_*phoA*_ reporter in the Δ*phoB* background, as *E. coli* Pho regulon genes have previously been shown to be induced by acidic conditions even in PhoB deletion mutants as well as in P replete conditions [35].

We observed that the average fold increase in fluorescence signal for the WT-background reporter strain was 6.1 for acidic pH, 6.6 for neutral pH, and 7.7 for alkaline pH (Figure 3A). Optical density measurements also indicate that the cells grew more poorly in the acidic condition compared to the neutral and alkaline conditions (Figure 3B), which may have contributed to the lower expression level observed in the acidic condition. The leaky mNG expression from the reporter, however, remained consistently low in replete P across all tested pH conditions. Furthermore, in the Δ*phoB* background, the reporter did not respond to P limitation under any of the tested pH conditions. These results together indicate that *P. synxantha* 2-79 does not appear to experience the acidity-induced Pho regulon upregulation that was observed in *E. coli* [35]. However, the fact that the expression level of the ON condition differs across pH highlights the importance of characterizing system performance across different pH values that are relevant to the soil environment.

**Figure 3:**
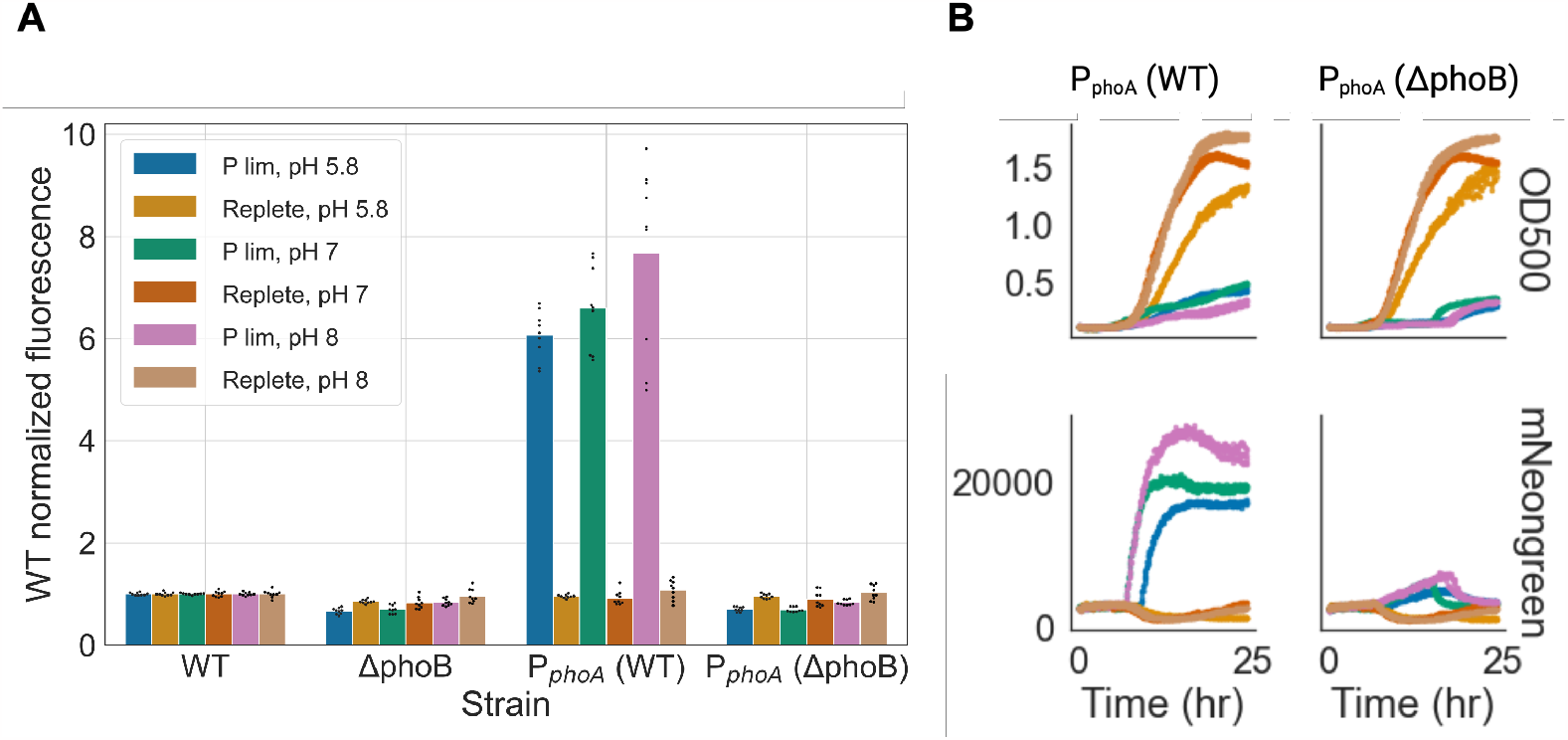
The P reporter strain in WT and ΔphoB background was grown in P limited and P replete media at three different pH conditions. A) Barplots of WT normalized fluorescence at 24 hours of growth. Bars show the average fluorescence value, dots show raw data for three biological replicates. B) Representative growth curves and mNG/OD for P_*phoA*_ in the WT and ΔphoB deletion strain.

### The reporter response is robust to the presence of organic P compounds

Up until this point, we have characterized the reporter’s behavior in low and high concentrations of inorganic P, which is accessible to plants. However, soil environments contain many other organic forms of P that are inaccessible as nutrients to plants but potentially accessible to bacteria [36] (Figure 4A). To properly indicate the limitation of bioavailable phosphorous, it is essential that our reporter be insensitive to concentrations of organic phosphorous.

**Figure 4:**
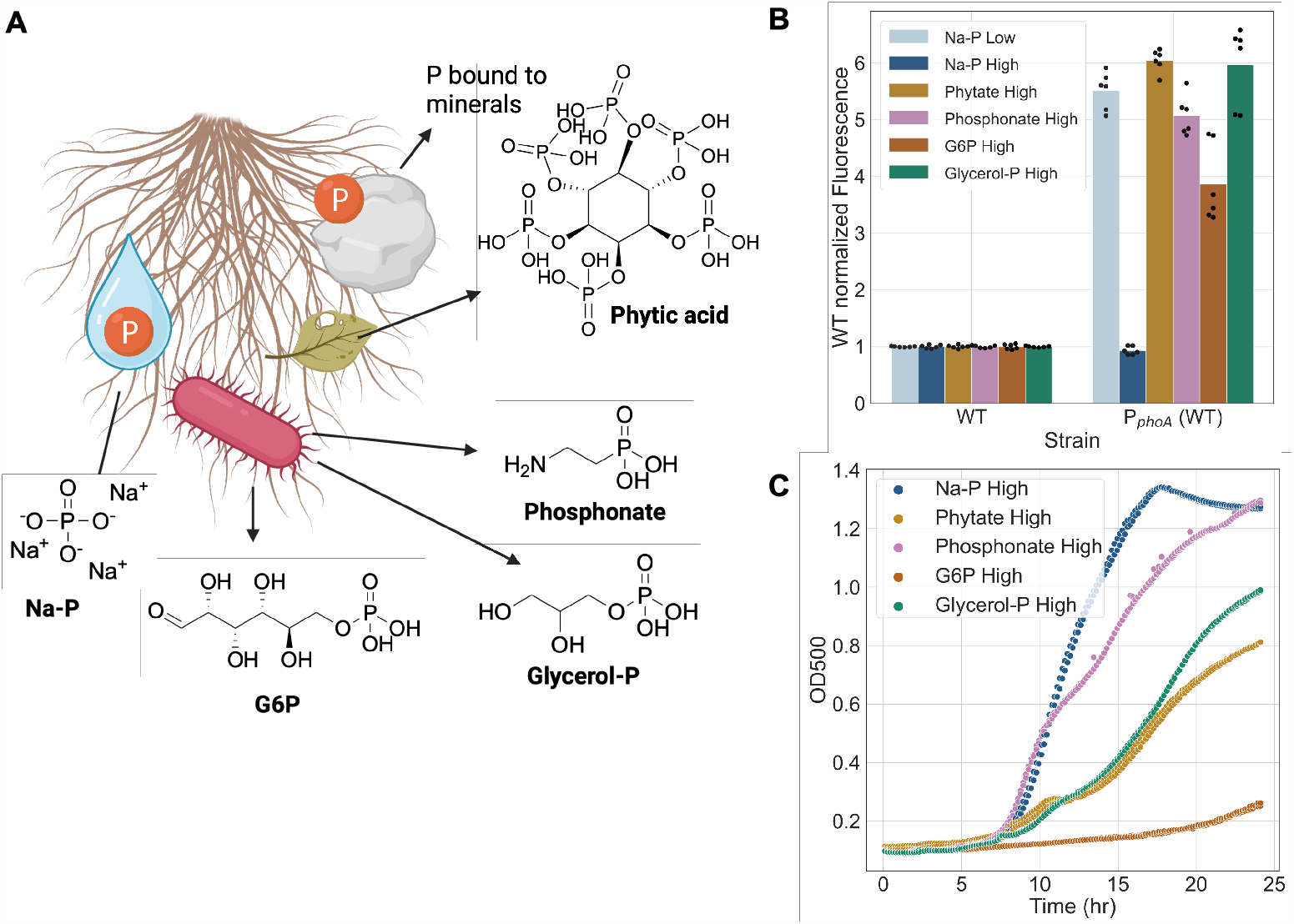
(A) Schematic showing examples of P types can be found as in the soil. Inorganic P can be tighly bound to minerals like calcium or iron or soluble in the pore water and available for plant uptake. Organic P in the soil can originate from dead plant material containing the main plant P storage molecule phytate. Organic P can also originate from other dead organisms like bacteria or it can come from anthropogenic sources, eg. in herbicides. Organic P cannot be utilized by plants for growth. (B) Reporter cells were grown in different P sources at replete conditions (1 mM). The WT normalized fluorescence is plotted at 24 hours of growth. Bars show the average fluorescence value, dots show raw data for three biological replicates. (C) Representative growth curves for one out of three biological replicates for the P replete condition.

To evaluate the robustness of our reporter’s response, we tested if the addition of organic P sources to a medium limited for inorganic P would rescue the reporter from the P limited condition and turn the reporter off. We predicted the output signal would remain on even after organic P is added to the medium if our reporter is insensitive to organic P sources. When we performed this experiment with four different organic P sources commonly found in soils, we found that none were able to reduce the reporter output down to replete levels (Figure 4B, Supplemental information: Figure S4). When limited for inorganic P (Na-P), the reporter exhibited a 5.5-fold increase in fluorescence over the WT background. When replete concentrations of Na-P (1 mM) were added, the reporter response went down to WT background levels, as expected. However, when the four organic P sources were added to 1 mM concentration, the reporter signal remained between 3.9- and 6-fold above WT levels. These results indicate that organic P sources are unable to rescue the reporter from its P limited state.

We note that some of the P sources are more difficult for the cells to import or degrade than others, which impacts the cells’ growth dynamics (Figure 4C). Phosphonate seems to be the most preferred organic P source, while the cells can barely utilize glucose 6-phosphate.

These results show that the P reporter signal is only silenced by inorganic P among the P sources tested in this experiment. This means that in the presence of phosphorus that is not available to plants, the limitation signal is on.

### Selection of a constitutive promoter for the ratiometric reporter

Having demonstrated that the P_*phoA*_ promoter can act as a specific and reliable indicator of inorganic P limitation in *P. synxantha* 2-79, we proceeded to develop our strain into a ratiometric bioreporter by incorporating the constitutive expression of a distinct fluorescent protein. Ratiometric readouts expand the utility of bioreporters by providing a measure of global cellular activity that can be used to calibrate the measured expression level from the inducible promoter. In environments like soils where only a fraction of the population may be active at any given time, such calibrations are essential for properly interpreting the reporter’s behavior. The constitutive expression signal can also be used to identify the spatial location of the bioreporters.

We chose to use the bright red fluorescent protein mScarletI [37] (RFP) as the constitutive reporter. In selecting the strength of the RFP output, it is important that it is strong enough to be easily detectable without being so strong that it sequesters cellular resources away from the expression of the output GFP signal [38]. We created three candidate dual reporter constructs where mNeonGreen is driven by P_*phoA*_ and mScarletI is driven by either the constitutive lac derived promoter Pa10403 [39] or by one of two synthetic Anderson promoters [40] that were previously found to express in *P. synxantha* [41]. We integrated these reporter constructs into the genome and measured the expression strengths of both fluorescent proteins under P limited and P replete conditions.

We observed that all three dual reporters exhibited a lower maximal GFP signal compared to the GFP-only reporter, which had a 5.9-fold increase in fluorescence over WT in the P limited condition (Figure 5A). Although the Pa10403 variant reduced this maximal GFP expression to 3.4-fold over WT, the two Anderson variants maintained GFP expression at 5-to 5.1-fold above WT. Interestingly, the RFP signal was higher in the P limited conditions than the P replete conditions for all three constitutive promoters (Figure 5B). Consistent with previous reports [41], the P1m promoter was stronger than the P2d promoter. We therefore chose to express the RFP from the P1m promoter in our final dual reporter construct, as it decreased GFP expression to a similar level as P2d while having a 2.8-fold higher RFP expression level.

**Figure 5:**
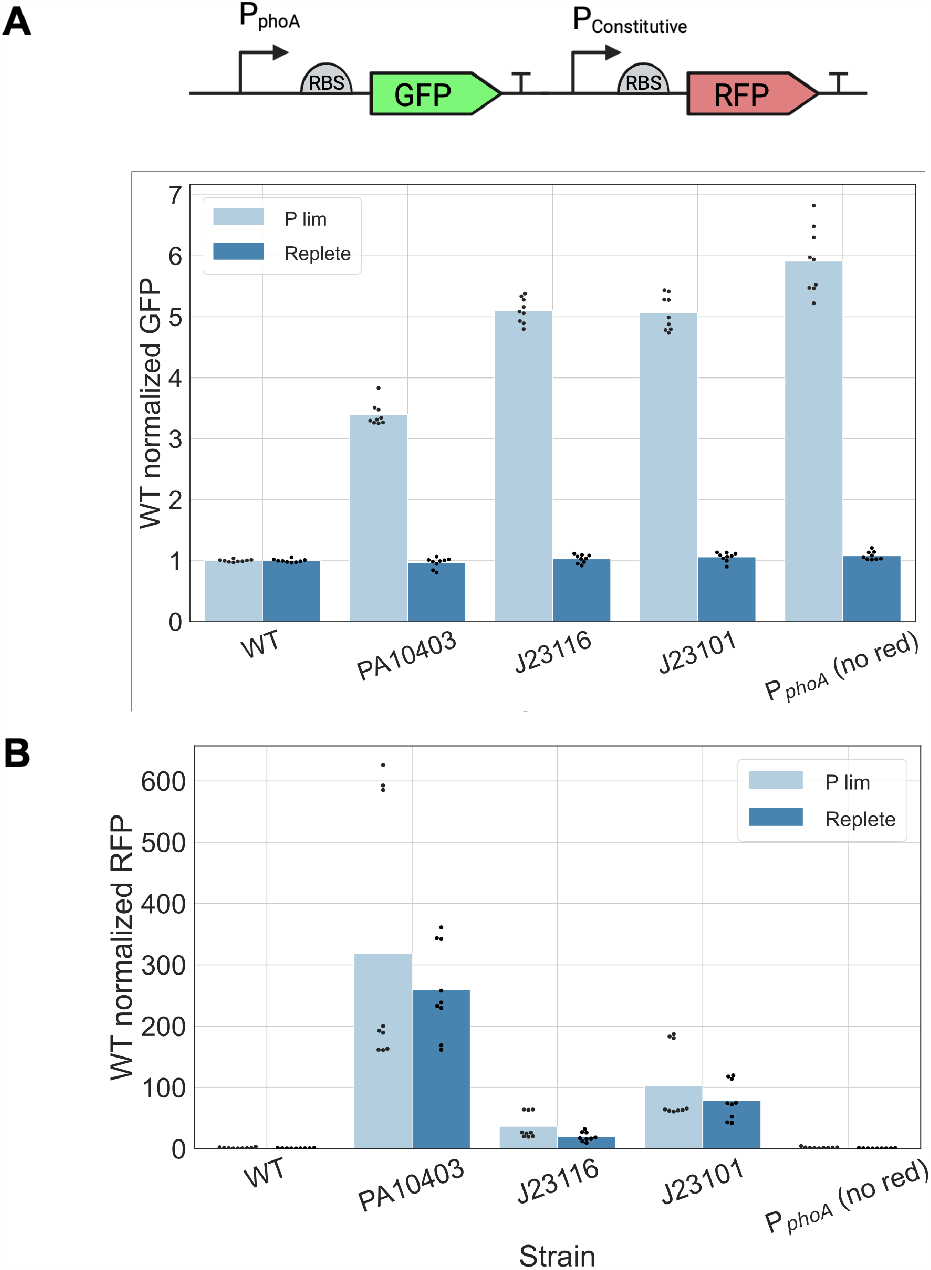
Three ratiometric variants were constructed and tested under P limited and replete conditions. (A) Comparison in GFP levels, normalized by WT, with three different constitutive promoters driving expression of mScarletI, compared to no RFP expression. (B) WT normalized RFP expression under three constitutive promoters compared to background fluorescence. Bars show the average fluorescence value, dots show raw data for three biological replicates.

### Ratiometric reporter performance in a soil context

To assess the performance of our dual reporter in a soil context, we grew the reporter strain in soil slurries that were generated by mixing soil from the Cook Agronomy Farm (CAF), from which *P. synxantha* 2-79 was isolated [42], with medium that was either limited (50 *µ*M) or replete (7 mM) with P. To verify soil slurries were limited for P, we measured aqueous P in abiotic controls after incubations for 24 hours. There was no detectable P present in these samples, indicating P in the limited medium had sorbed to the soil (Supplemental information C, Table S2). Similarly, lower soluble P was measured in soil samples that had been mixed with replete P medium.

Cells were grown for 24h in the slurry and then extracted for analysis by flow cytometry (Figure 6A, Supplemental information C). As a comparison, we also analyzed the response of the reporter grown in pure growth medium to determine whether it is predictive of the reporter’s performance in the soil context.

**Figure 6:**
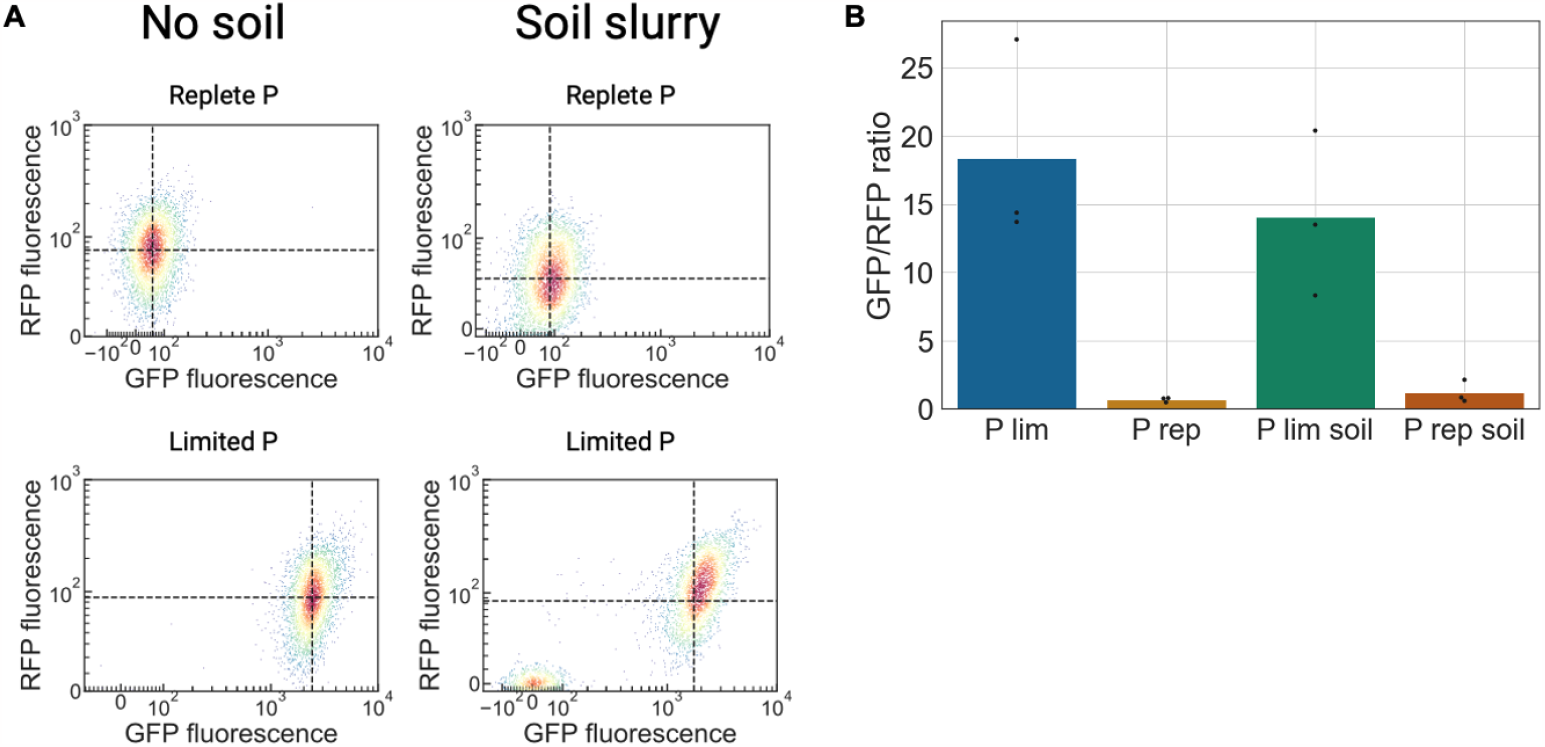
Flow cytometry measurement of three biological replicates taken on three different days in liquid cultures and soil slurries.(A) Representative plot of density gated flow cytometry data for one out of three biological replicates. The median intensities of GFP and RFP fluorescence are identified at the intersection of the dashed lines. (B) The median fluorescence intensities for GFP and RFP are divided by each other for each biological replicate. Bars show the average ratio, dots show each ratio for three biological replicates.

The use of calibration beads to measure fluorescent protein expression in absolute fluorescent units enabled us to directly compare the measured values across different conditions. Expression strengths of both GFP and RFP were generally similar between the soil slurry and pure growth medium within a P condition (Figure 6A). The major distinction came from the fact that 29-64% of the cells in the P limited soil slurry had minimal expression levels of both GFP and RFP, suggesting that they are metabolically inactive or dead.

We then computed the median GFP and RFP fluorescence values for each condition and divided them to obtain a GFP/RFP ratio that corresponds to the ratiometric readout value for each condition (Fig 6B). We can observe that in the soil slurry, the reporter has a higher leak (1.2 compared to 0.7) and lower dynamic range than in the pure growth medium. However, the reporter nonetheless maintains a 12-fold response to the limitation of P in the soil slurry.

Taken together, these results indicate that soil context affects the performance of the reporter in two major ways. First, it creates an inactive fraction of the population. Second, it reduces the dynamic range by both increasing the leaky expression level and decreasing the maximal expression level. However, despite these effects, our ratiometric reporter can still provide a reliable readout for P limitation in the soil slurry with a 12-fold dynamic range. Given that the reporter exhibits a 27-fold dynamic range in pure growth medium, further improvements need to be made to increase performance robustness in the soil environment. Future experiments to better understand the physiological impacts of the soil context will help expand the applicability of the reporter to different types of environmental contexts.

## Conclusion

This study presents a framework for testing environmentally relevant parameters when engineering a bacterial bioreporter in a bacterium isolated from the context of its intended application, in this case, the wheat rhizosphere of the Columbia Plateau. Our work demonstrates the utility of selecting a non-native promoter for synthetic biology applications in a new chassis organism, and characterizing its response under environmentally relevant conditions (pH, P sources, soil).

Future steps to refine the reporter for rhizosphere applications include further characterization of the ratiometric reporter in conditions more similar to the actual soil environment, for example by reducing the water content or examining its performance in the presence of native soil microbes by not autoclaving the soil prior to the experiments. Another worthwhile pursuit would be to alter the promoter region to achieve stronger GFP expression.

Although long-term *in situ* use of genetically engineered bioreporters is not practical today, work towards understanding what underpins lonegvity and reliable performance over time is necessary for bioreporters to realize their full potential in agricultural applications. The modular structure of many bioreporters can be leveraged both to sense other parameters of interest (e.g. different nutrients, such as ammonium or nitrate) and/or to enable actuation rather than reporting, opening up the possiblity of modulating the soil environment for bioremediation or liberation of nutrients bound to minerals.

## Methods

### Construction and genome integration of *P. synxantha* reporters

All cloning to produce *P. synxantha* genomic integration constructs was done using *E. coli* DH10B (Invitrogen) with the backbone pJM220 [43]. For the native promoters the intergenic region upstream of the *pstS* (locus tag C4K02 RS29030) and *phoX* (locus tag C4K02 RS26910) genes were amplified via PCR adding Gibson overhangs for the pJM220 backbone vector. Gibson assembly was then done for each construct and transformed into *E. coli* DH10B competent cells. Plasmids were purified using a QIAprep Spin Miniprep kit (Qiagen).

The constructs were then integrated on the *P. synxantha* chromosome using transposase based insertion at the Tn7 site. The protocol used for making and transforming competent cells was modified from Choi et al. [23].

Briefly, electrocompetent *P. synxantha* cells were electroporated in 1 mm-gap cuvettes (at 1.8 mV, 600 Ω and 10 *µ*F) with the construct plasmid as well as a plasmid containing the transposase and genes required for genome insertion [44]. The cells were then recovered in rich medium (SOC) for 3 hours at 30°C and plated onto LB agar plates containing gentamicin (20 *µ*g/ml) and incubated for 24 hours before picking colonies for sequence verification.

### Plate reader assay for *in vivo* fluorescence

*In vivo* fluorescence was measured using a Biotek (Synergy H1) plate reader. The experiments ran for 24 hours at 30°C using continuous orbital shaking starting from an overnight culture diluted to OD 0.1. OD was measured every 10 minutes at 500 nm and fluorescence was measured at 490/520 nm for mNeonGreen and at 569/593 nm for mScarletI. In experiments where phenazine-1-carboxylic acid was assessed, absorbance at 367 nm was measured. The background fluorescence of the wild-type strain was subtracted from the fluorescence values of the reporter strains in Figure 2B.

### Measurement of phosphate supernatant concentration

Measurements of phosphate concentration in the growth medium supernatant were done using a Malachite green phosphate assay kit (Sigma-Aldrich, Cat. No. MAK307). After collecting the supernatant, the sample was filtered through a 0.22 *µ*m filter and stored at -20C.

### Preparation of cells for flow cytometry

Starting from an individual colony, a culture was grown in LB medium overnight. 1 ml of the culture was pelleted and washed once in minimal P limited medium. The cells were then resuspended in 1 ml minimal P limited medium (30 uM P). For the no soil condition, 3 ml of minimal medium (P limited or replete) was inoculated with the washed overnight culture (at OD 0.15). For the soil condition, tubes filled with 4 g of autoclaved soil were filled with 4 ml medium (P limited or replete). The tubes were then vortexed to mix the soil and liquid to yield a slurry. The tubes were then inoculated at the same cell density as the no soil cultures. All cultures were grown for 24 hours at 30°C and 220 rpm shaking. The soil in the tubes was then separated from the bacteria following the protocol in Chemla et al. [9]. All cultures were then diluted 1:100 and filtered through a 10 *µ*m filter before proceeding to flow cytometry measurements.

### Flow cytometry and data analysis

The flow cytometry measurements were performed on a CytoFLEX S flow cytometer using the 610 Yellow and 525 Blue lasers. Approximately 10000 events were collected for each sample. In each experiment, the fluorescence of calibration beads (Spherotech) was measured to enable convertion of the arbitrary fluorescence units (AFU) to molecules of equivalent fluorophore (MEF) using the pipeline developed by Castillo-Hair et al. [45].

The analysis followed the Castillo-Hair et al. protocol [45]. First, the calibration bead data was used to convert the arbitrary units on the machine to MEF. Then, background noise that was not single cells was gated out and 40 percent of the events in the densest region were kept for further analysis. From the remaining events, the median fluorescence was calculated for both flourescent proteins. The GFP values were then divided by the RFP values for each experimental condition.

### Media recipes

For routine growth we used LB and LB agar. For limitation assays, the cells were grown in minimal media containing 0.41 mM MgSO4, 0.68 mM CaCl2 and 25 mM MOPS (or 12.5 mM MES for low pH medium). Aquil trace metals [46] were added containing 10 *µ*M Fe and 100 *µ*M EDTA. The limited media were designed according to the Redfield ratio with guidance from D. McRose [47]. For P limited medium, the final concentration of potassium phosphate was 20, 30, 50 or 100 *µ*M. For N limited medium, the final concentration of ammonium chloride was 1 mM. For C limited medium the final concentration of glucose was 6.625 mM.

In the P source experiment, the concentration of P source (sodium phosphate, 2-aminoethyl phosponic acid, phytic acid, glucose 6-phosphate or glycerol 3-phosphate) in the replete condition was 1 mM and the limited condition was 50 *µ*M. The concentration of added KCl was 1 mM in both limited and replete media.

## Author information

Project idea - DKN and EML

Experimental design - EML, RMM and DKN

Experiments/data collection - EML except one of the four malachite green assay measurements that was performed by DKN.

Data analysis and visualization - EML

Writing manuscript - EML

Editing of manuscript - EML, RMM and DKN

## Acknowledgements

We thank R. Alcalde for insightful discussions regarding project conceptualization and for creating the P_*phoA*_-mNeonGreen strain. We thank I. Antoshechkin at the Millard and Muriel Jacobs Genetics and Genomics Laboratory at Caltech for assistance in pre-processing the RNA sequencing raw data. We also thank D. McRose for sharing the initial Δ*phoB* strain, and thank J. Marken, D. Mavrodi, O. Mavrodi, G. Squyres, L. Tsypin, and S. Wilbert for fruitful discussions that helped move this work forward. All figures were created using BioRender.com. This work was supported by the Institute for Collaborative Biotechnologies through contract W911NF-19-D-0001 from the U.S. Army Research Office and Caltech’s Resnick Sustainability Institute. The content of the information in this report does not necessarily reflect the position or the policy of the Government, and no official endorsement should be inferred.

## References

[1] A. Babapoor, R. Hajimohammadi, S. M. Jokar, and M. Paar, “Biosensor design for detection of mercury in contaminated soil using rhamnolipid biosurfactant and luminescent bacteria,” Journal of Chemistry, vol. 2020, pp. 1–8, 2020.

[2] Y. Wu, C.-W. Wang, D. Wang, and N. Wei, “A whole-cell biosensor for point-of-care detection of waterborne bacterial pathogens,” ACS Synthetic Biology, vol. 10, no. 2, pp. 333–344, 2021.

[3] K. M. DeAngelis, P. Ji, M. K. Firestone, and S. E. Lindow, “Two novel bacterial biosensors for detection of nitrate availability in the rhizosphere,” Applied and Environmental Microbiology, vol. 71, no. 12, pp. 8537–8547, 2005.

[4] R. Gebbers and V. I. Adamchuk, “Precision agriculture and food security,” Science, vol. 327, no. 5967, pp. 828–831, 2010.

[5] D. L. Jones and E. Oburger, “Solubilization of phosphorus by soil microorganisms,” Phosphorus in action: biological processes in soil phosphorus cycling, pp. 169–198, 2011.

[6] G. Recorbet, C. Picard, P. Normand, and P. Simonet, “Kinetics of the persistence of chromosomal DNA from genetically engineered Escherichia coli introduced into soil,” Applied and Environmental Microbiology, vol. 59, no. 12, pp. 4289–4294, 1993.

[7] L. S. Thomashow and D. M. Weller, “Role of a phenazine antibiotic from Pseudomonas fluorescens in biological control of Gaeumannomyces graminis var. tritici,” J. Bacteriol., vol. 170, pp. 3499–3508, Aug. 1988.

[8] Y. Chen, X. Shen, H. Peng, H. Hu, W. Wang, and X. Zhang, “Comparative genomic analysis and phenazine production of Pseudomonas chlororaphis, a plant growth-promoting rhizobacterium,” Genom Data, vol. 4, pp. 33–42, June 2015.

[9] Y. Chemla, Y. Dorfan, A. Yannai, D. Meng, P. Cao, S. Glaven, D. B. Gordon, J. Elbaz, and C. A. Voigt, “Parallel engineering of environmental bacteria and performance over years under jungle-simulated conditions,” PloS One, vol. 17, no. 12, p. e0278471, 2022.

[10] L. De Weger, L. Dekkers, A. Van Der Bij, and B. J. Lugtenberg, “Use of phosphate-reporter bacteria to study phosphate limitation in the rhizosphere and in bulk soil,” Molecular Plant-Microbe interactions, vol. 7, no. 1, pp. 32–38, 1994.

[11] L. Kragelund, C. Hosbond, and O. Nybroe, “Distribution of metabolic activity and phosphate starvation response of lux-tagged Pseudomonas fluorescens reporter bacteria in the barley rhizosphere,” Applied and Environmental Microbiology, vol. 63, no. 12, pp. 4920–4928, 1997.

[12] M. A. Dollard and P. Billard, “Whole-cell bacterial sensors for the monitoring of phosphate bioavailability,” J. Microbiol. Methods, vol. 55, pp. 221–229, Oct. 2003.

[13] E. Dekel and U. Alon, “Optimality and evolutionary tuning of the expression level of a protein,” Nature, vol. 436, no. 7050, pp. 588–592, 2005.

[14] L. López-Maury, S. Marguerat, and J. Bähler, “Tuning gene expression to changing environments: from rapid responses to evolutionary adaptation,” Nature Reviews Genetics, vol. 9, no. 8, pp. 583–593, 2008.

[15] W. R. Whitaker, E. S. Shepherd, and J. L. Sonnenburg, “Tunable expression tools enable single-cell strain distinction in the gut microbiome,” Cell, vol. 169, no. 3, pp. 538–546, 2017.

[16] K. Temme, R. Hill, T. H. Segall-Shapiro, F. Moser, and C. A. Voigt, “Modular control of multiple pathways using engineered orthogonal T7 polymerases,” Nucleic Acids Research, vol. 40, no. 17, pp. 8773–8781, 2012.

[17] R. D. Monds, P. D. Newell, J. A. Schwartzman, and G. A. O’Toole, “Conservation of the Pho regulon in Pseudomonas fluorescens Pf0-1,” Appl. Environ. Microbiol., vol. 72, pp. 1910–1924, Mar. 2006.

[18] S. G. Gardner and W. R. McCleary, “Control of the phoBR regulon in Escherichia coli,” EcoSal Plus, vol. 8, no. 2, pp. 10–1128, 2019.

[19] F. Santos-Beneit, “The Pho regulon: a huge regulatory network in bacteria,” Front. Microbiol., vol. 0, 2015.

[20] B. Wanner, “Gene regulation by phosphate in enteric bacteria,” Journal of Cellular Biochemistry, vol. 51, no. 1, pp. 47–54, 1993.

[21] N. C. Shaner, G. G. Lambert, A. Chammas, Y. Ni, P. J. Cranfill, M. A. Baird, B. R. Sell, J. R. Allen, R. N. Day, M. Israelsson, et al., “A bright monomeric green fluorescent protein derived from branchiostoma lanceolatum,” Nature Methods, vol. 10, no. 5, pp. 407–409, 2013.

[22] M. B. Elowitz and S. Leibler, “A synthetic oscillatory network of transcriptional regulators,” Nature, vol. 403, no. 6767, pp. 335–338, 2000.

[23] K.-H. Choi and H. P. Schweizer, “mini-Tn7 insertion in bacteria with single att Tn7 sites: example Pseudomonas aeruginosa,” Nature Protocols, vol. 1, no. 1, pp. 153–161, 2006.

[24] D. V. Mavrodi, V. N. Ksenzenko, R. F. Bonsall, R. J. Cook, A. M. Boronin, and L. S. Thomashow, “A seven-gene locus for synthesis of phenazine-1-carboxylic acid by Pseudomonas fluorescens 2-79,” Journal of Bacteriology, vol. 180, no. 9, pp. 2541–2548, 1998.

[25] W. S. Kisaalita, P. J. Slininger, and R. J. Bothast, “Defined media for optimal pyoverdine production by Pseudomonas fluorescens 2-79,” Applied Microbiology and Biotechnology, vol. 39, pp. 750–755, 1993.

[26] H. M. Bilal, T. Aziz, M. A. Maqsood, M. Farooq, and G. Yan, “Categorization of wheat genotypes for phosphorus efficiency,” PloS One, vol. 13, no. 10, p. e0205471, 2018.

[27] A. Puri-Taneja, S. Paul, Y. Chen, and F. M. Hulett, “Ccpa causes repression of the phopr promoter through a novel transcription start site, pa6,” Journal of Bacteriology, vol. 188, no. 4, pp. 1266–1278, 2006.

[28] J. F. Martín, A. Sola-Landa, F. Santos-Beneit, L. T. Fernández-Martínez, C. Prieto, and A. Rodríguez-García, “Cross-talk of global nutritional regulators in the control of primary and secondary metabolism in streptomyces,” Microbial Biotechnology, vol. 4, no. 2, pp. 165–174, 2011.

[29] P. Bielecki, V. Jensen, W. Schulze, J. Gödeke, J. Strehmel, D. Eckweiler, T. Nicolai, A. Bielecka, T. Wille, R. G. Gerlach, et al., “Cross talk between the response regulators PhoB and TctD allows for the integration of diverse environmental signals in Pseudomonas aeruginosa,” Nucleic Acids Research, vol. 43, no. 13, pp. 6413–6425, 2015.

[30] F. Demoling, D. Figueroa, and E. Bååth, “Comparison of factors limiting bacterial growth in different soils,” Soil Biology and Biochemistry, vol. 39, no. 10, pp. 2485–2495, 2007.

[31] S. Heinze, J. Raupp, and R. G. Joergensen, “Effects of fertilizer and spatial heterogeneity in soil ph on microbial biomass indices in a long-term field trial of organic agriculture,” Plant and Soil, vol. 328, pp. 203–215, 2010.

[32] R. Smiley and R. Cook, “Relationship between take-all of wheat and rhizosphere ph,” Phytopathology, vol. 63, pp. 882–890, 1973.

[33] R. Smiley, “Rhizosphere pH as influenced by plants, soils, and nitrogen fertilizers,” Soil Science Society of America Journal, vol. 38, no. 5, pp. 795–799, 1974.

[34] A. Ortega-Pieck, J. Norby, E. S. Brooks, D. Strawn, A. R. Crump, and D. R. Huggins, “Sources and subsurface transport of dissolved reactive phosphorus in a semiarid, no-till catchment with complex topography,” tech. rep., Wiley Online Library, 2020. doi:10.1002/jeq2.20114.

[35] L. W. Marzan and K. Shimizu, “Metabolic regulation of Escherichia coli and its phoB and phoR genes knockout mutants under phosphate and nitrogen limitations as well as at acidic condition,” Microb. Cell Fact., vol. 10, p. 39, May 2011.

[36] H. Lambers, “Phosphorus acquisition and utilization in plants,” Annual Review of Plant Biology, vol. 73, pp. 17–42, 2022.

[37] D. S. Bindels, L. Haarbosch, L. Van Weeren, M. Postma, K. E. Wiese, M. Mastop, S. Aumonier, G. Gotthard, A. Royant, M. A. Hink, et al., “mScarlet: a bright monomeric red fluorescent protein for cellular imaging,” Nature Methods, vol. 14, no. 1, pp. 53–56, 2017.

[38] C. D. McBride, T. W. Grunberg, and D. Del Vecchio, “Design of genetic circuits that are robust to resource competition,” Current Opinion in Systems Biology, vol. 28, p. 100357, 2021.

[39] M. Lanzer and H. Bujard, “Promoters largely determine the efficiency of repressor action.,” Proceedings of the National Academy of Sciences, vol. 85, no. 23, pp. 8973–8977, 1988.

[40] “jPromoters/Catalog/Anderson - parts.igem.org — parts.igem.org.” http://parts.igem.org/Promoters/Catalog/Anderson. [Accessed 05-09-2023].

[41] J. T. Meyerowitz, E. M. Larsson, and R. M. Murray, “Development of cell-free transcription-translation systems in three soil pseudomonads,” bioRxiv, 2023. doi:10.1101/2023.06.09.544292.

[42] D. Weller, R. Cook, et al., “Suppression of take-all of wheat by seed treatments with fluorescent pseudomonads,” Phytopathology, vol. 73, no. 3, pp. 463–469, 1983.

[43] J. Meisner and J. B. Goldberg, “The Escherichia coli rhaSR-PrhaBAD inducible promoter system allows tightly controlled gene expression over a wide range in Pseudomonas aeruginosa,” Applied and Environmental Microbiology, vol. 82, no. 22, pp. 6715–6727, 2016.

[44] K.-H. Choi, J. B. Gaynor, K. G. White, C. Lopez, C. M. Bosio, R. R. Karkhoff-Schweizer, and H. P. Schweizer, “A Tn 7-based broad-range bacterial cloning and expression system,” Nature Methods, vol. 2, no. 6, pp. 443–448, 2005.

[45] S. M. Castillo-Hair, J. T. Sexton, B. P. Landry, E. J. Olson, O. A. Igoshin, and J. J. Tabor, “Flowcal: a user-friendly, open source software tool for automatically converting flow cytometry data from arbitrary to calibrated units,” ACS Synthetic Biology, vol. 5, no. 7, pp. 774–780, 2016.

[46] R. A. Andersen, Algal culturing techniques. Elsevier, 2005.

[47] D. L. McRose and D. K. Newman, “Redox-active antibiotics enhance phosphorus bioavailability,” Science, vol. 371, pp. 1033–1037, Mar. 2021.

