## Supplemental Methods, Tables and Figures for "Engineering the soil bacterium *Pseudomonas synxantha* 2-79 into a ratiometric bioreporter for phosphorus limitation"

Supplemental information

**A – P reporter design, construction, and screening**

Table S1

| **Promoter name** | **Sequence** |
| --- | --- |
| P_phoA_ (*E. coli*) | ATGAGCTCAAAAGTTAATCTTTTCAACAG**CTGTCAT**AAAG**TTGTCAC**GGCCGAGACTTATAGTCGCTTTGTTTTTATTTTTTAA |
| P_pstS_ (RS29030) | GCTAAAGCCAGCCTGATCCACTGTGGCGAGGGAGCTTGCTCCCGCTGGGGCGCGAAGCGGCCCTTAAAGATGGGACTGCTGCGCAGTCCAACGGGAGCAAGCTCCCTCGCCACAATGAAAGTGTTTGTTCGGTGTTTTACCTCGCATACCTTCGCTTCCCGATAGTGGC**CTTTCAT**CTAA**TTGTCAT**ATTTA**AG**A**CAT**AGAGTGTTCACACGGCCTCCCGATACTTGGCCCCGATCCAATTATCCCTATCTGCTAGGAGCAAGGC |
| P_acpA_ (RS03720) | TCACCTCGGTGATGTGCATTAGAAACTCAGTCTAGCAGCGCTTACTCTGTGTAGACACGCCCCACCCATCACTTGTGTGTCTTCGTTCCGTCACATTCCATTGCTACCGTCCTGCGAAATATCTTGCAACATCTCGCCAAGGACTCCCCCGCAAT |
| P_pstS_ (RS14750) | GGCGCTGGCGGATCCGAGCATTTTAGTCGCATTTTGTTACTAACCAATGCACTAAAAATATAGGGCTGTAGTGGCAAACCGGGACAAAACCCCGGTTTGCCACTCAAT**TTTCAT**AAAAATGTAACAACCCTGCTGCACAGTTCGGGAGCCCGTCACCACAACGAGTTCCTTTTA |
| P_phoX_ (RS26910)  Monds et al [1] | TGGGCGAACGTCTTCAAA**CTGCAA**CAATCTG**GTCA**CAAGTGCTGCTTAAGGTGT**CTGCAA**AAACCGCAGGGAGCCTGAG |
| P_phoD_ (RS04350)  Monds et al [1] | GTGTTGACGGGGTTGGTGTATGCGCTGCTTAAGCGACCGGAAGCTGTGGAACTGACGGTCACTGCTGCCAAGGGCTGATGGGATTCAAAATGTGGGAGGGGGCTTGCCCCCGATAGCGATGTATCAGTCAGCCTATTTTTAGCTGACACACTGCTATCGGGAGCAAGCCCCCTCCCACATTTGGATTTTCATTGGCTGTAAGGGTG**GTGTCA**TCTCTGC**TTCAT**CCCTACGTGCTTAAGTGGCCCTTTTGATACTCAAGAGGGCACCTGC |

Material and Methods: RNA sequencing (sample collection, RNA extraction, data preprocessing)

Cells (wild-type and the *ΔphoB* mutant) were grown from a single colony in minimal medium overnight. The cells were then washed three times and diluted to OD 0.05 into P limited medium at 35 μM P and grown to an OD of ~0.3. The cells were then separated into two 15 ml tubes at a volume of 6 ml each and P was added to one of the tubes to reach a P concentration of 5 mM and incubated at 30C for 5 minutes. The cell cultures were then centrifuged at 6200 xg and the supernatant was decanted, and cell pellets were immediately flash frozen by submerging the tubes into liquid nitrogen. Cell pellets were stored at -80C until performing RNA extractions.

A Qiagen RNeasy kit was used for the RNA extractions and RNA concentrations were determined using a Nanodrop. RNA sequencing was done at the Millard and Muriel Jacobs Genetics and Genomics Laboratory at Caltech.

The raw data was pre-processed using STARaligner, featureCounts and DESeq. Sorting on lowest p-value, the three most upregulated genes, that were not also upregulated in the *ΔphoB* mutant, were selected as candidates for P reporter constructs.

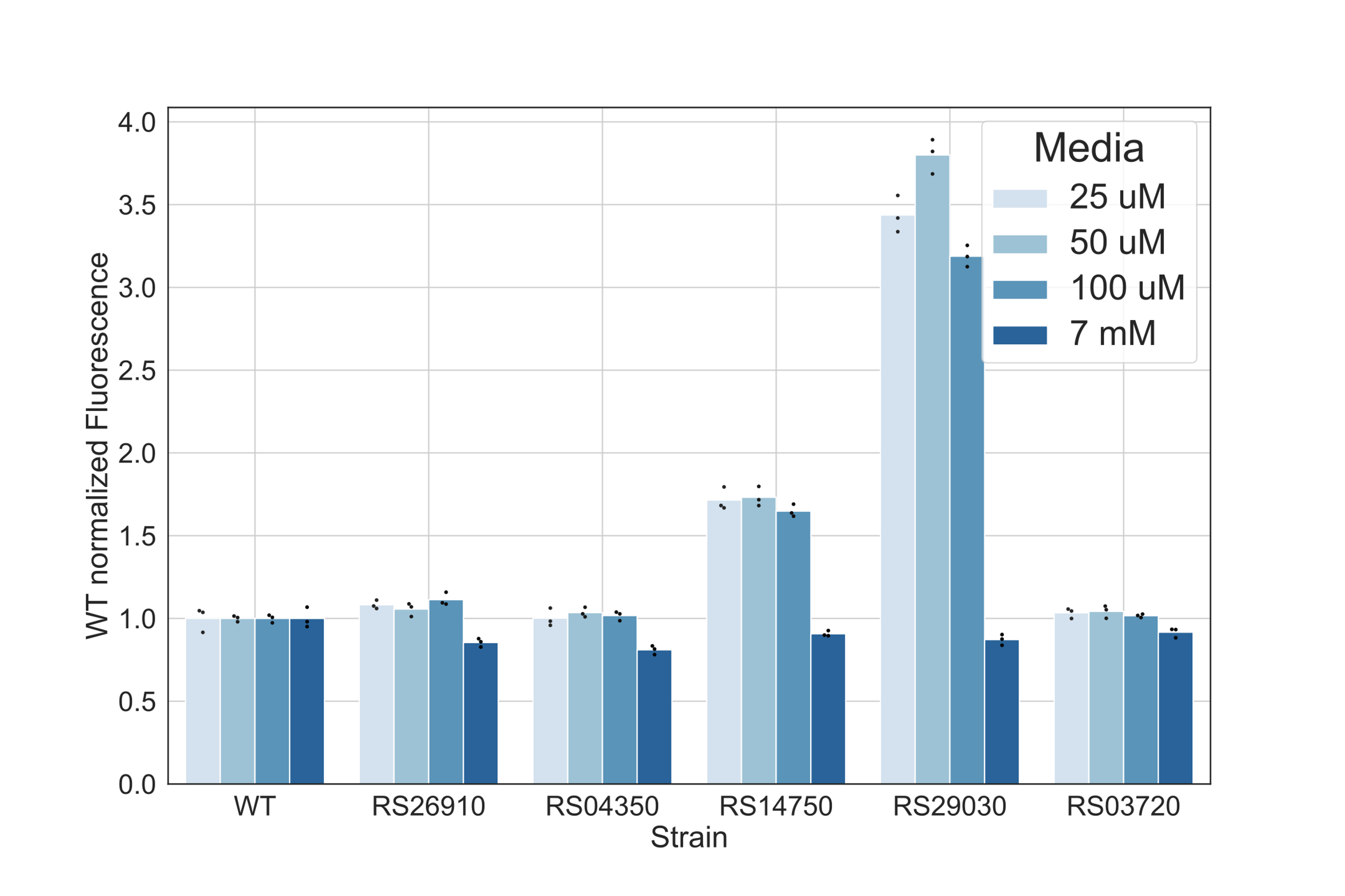

**Figure S1:** Screen of promoter fusions for 5 native *P. synxantha* intergenic regions. Promoter fusion strains were grown in minimal media with different concentrations of phosphorus for 24 hours. End point GFP fluorescence is normalized by the fluorescence of WT cells without any integrated promoter fusion.

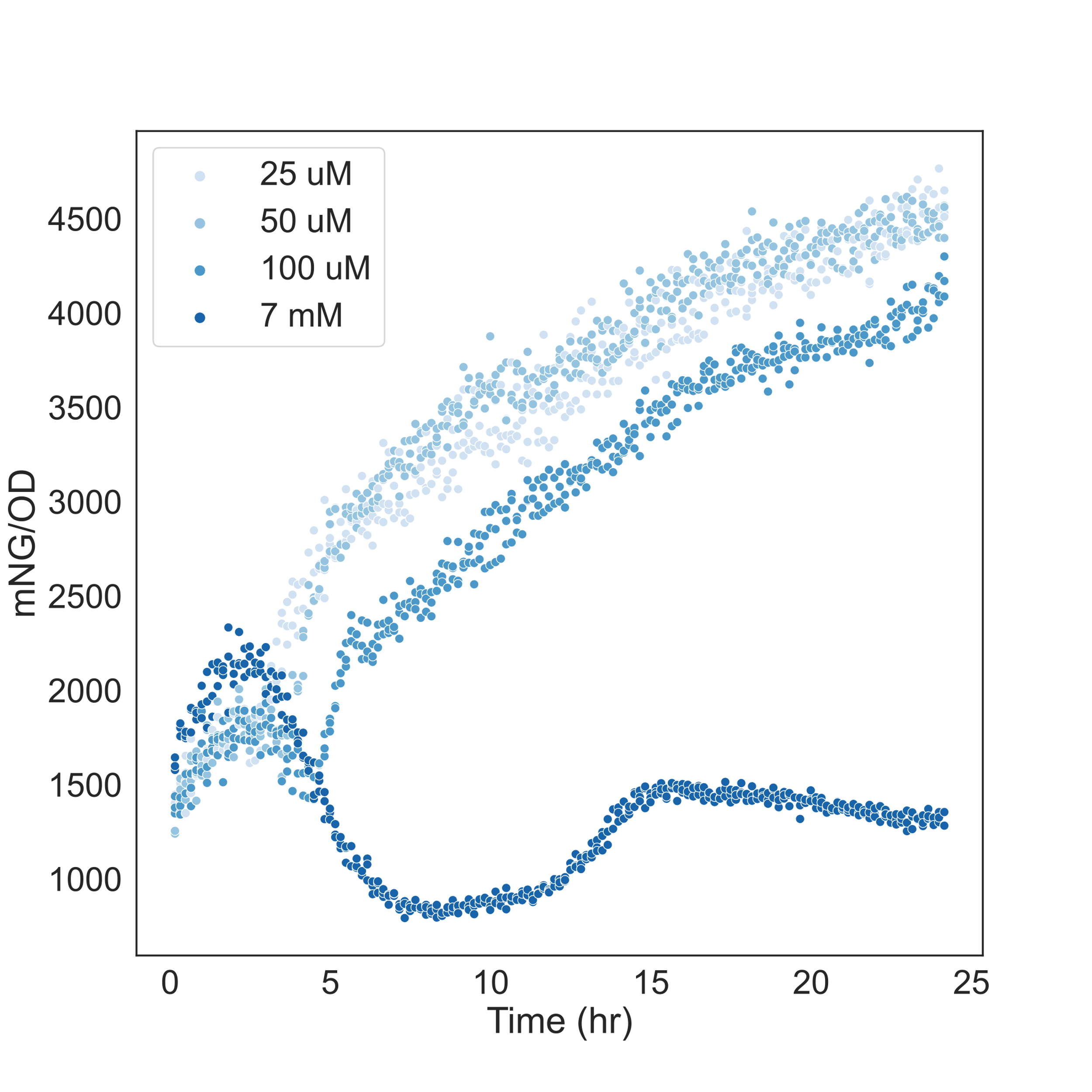

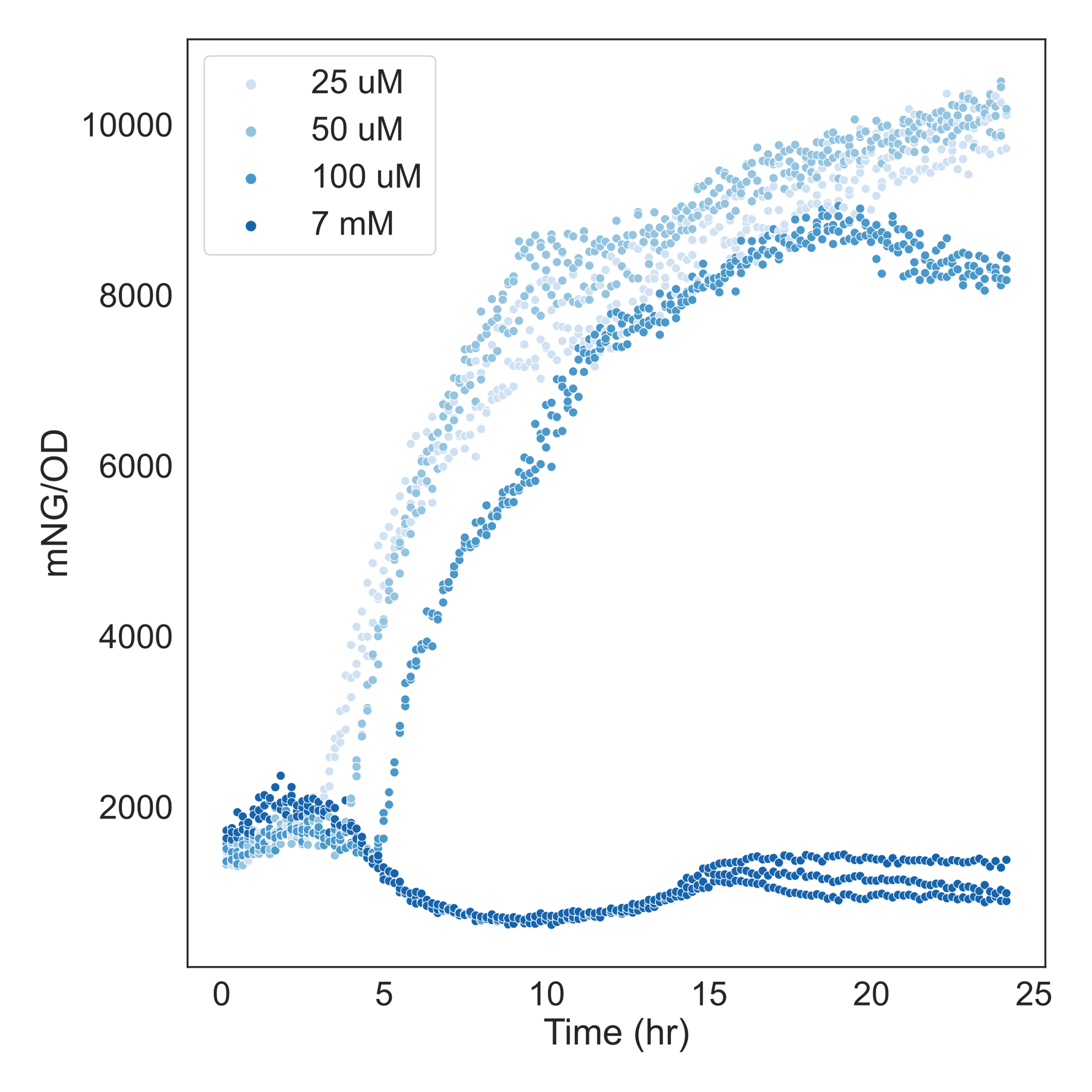

**Figure S2:** mNG fluorescence normalized by OD for 24 hours of growth. Upper: Promoter RS14750. Lower: Promoter RS29030.

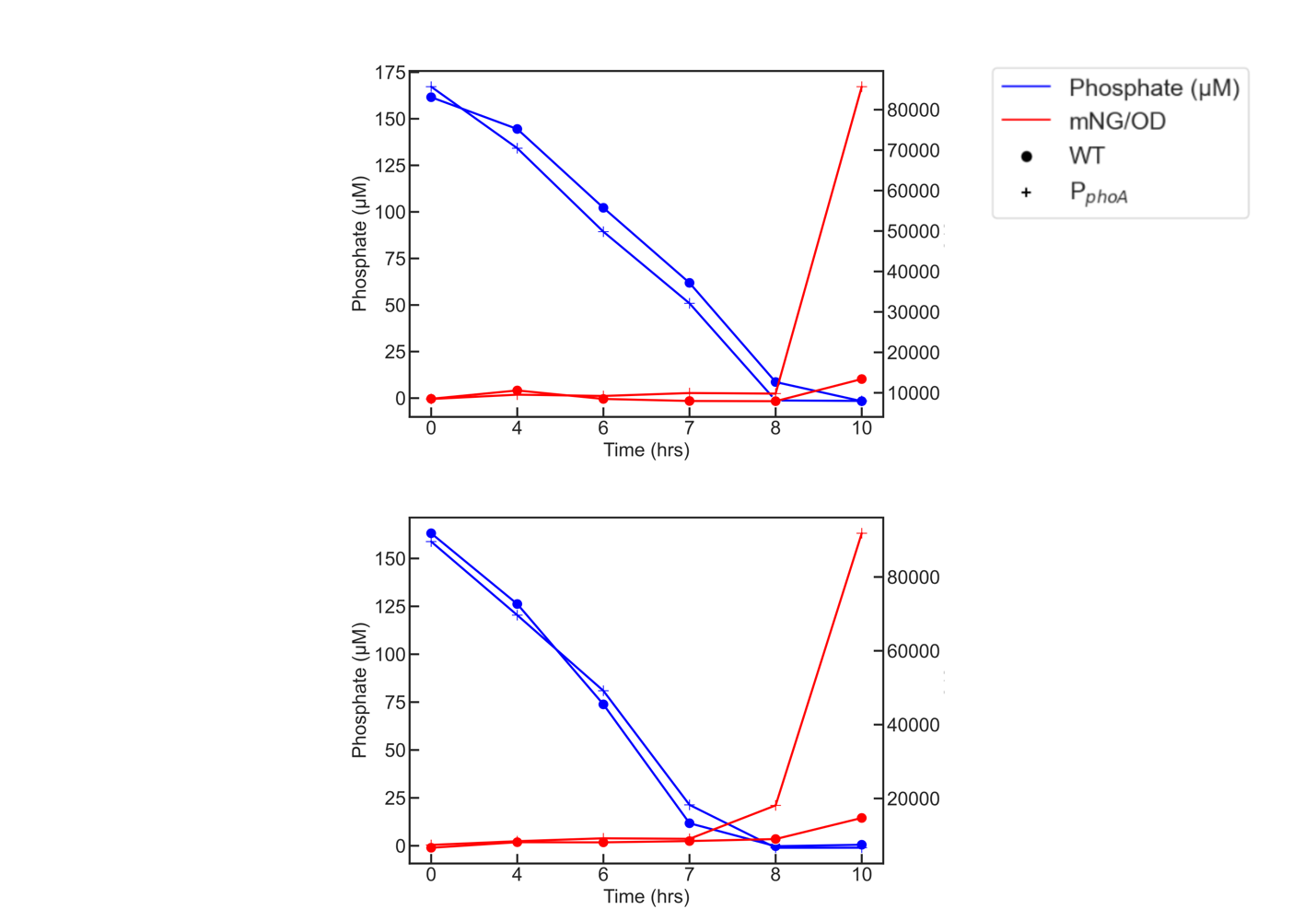

**Figure S3:** Phosphate, OD and fluorescence measurements over 10 hours of growth of WT and P_phoA_ cells. Biological replicates from two separate days.

**B – P source experiment**

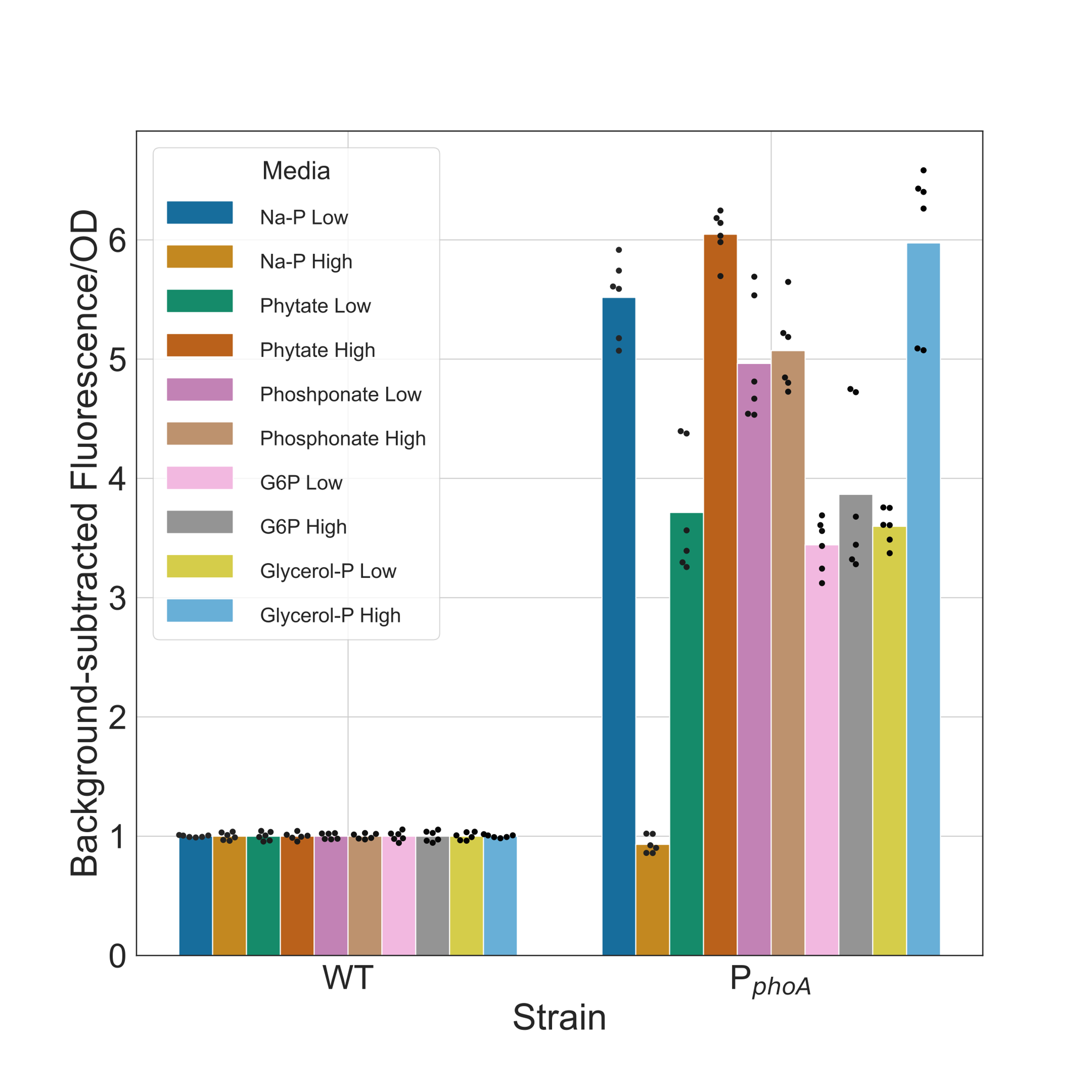

**Figure S4:** End point fluorescence after 24 hours of growth in different P sources. Low indicates 50 uM and High indicates 1 mM of added P.

**C – Soil slurry flow cytometry**

Table S2: Malachite green assay P concentrations after 24 h incubation

| **Sample** | **[Soluble P]** |
| --- | --- |
| Limited medium (no soil) | 35 μM |
| Limited medium + soil (replicate #1) | Not detectable (nd) |
| Limited medium + soil (replicate #2) | nd |
| Limited medium + soil (replicate #3) | nd |
| Replete medium + soil | 2300 μM |

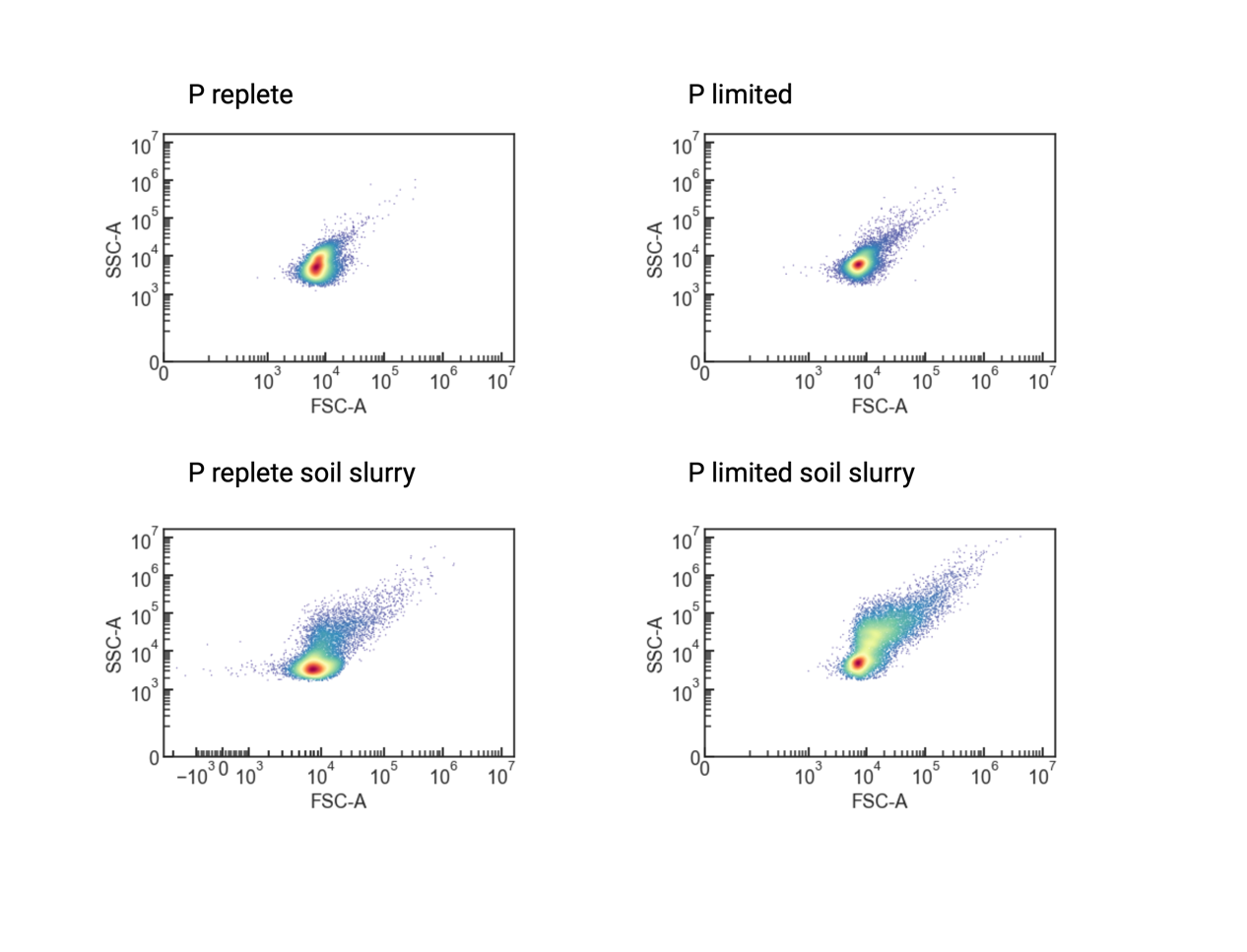

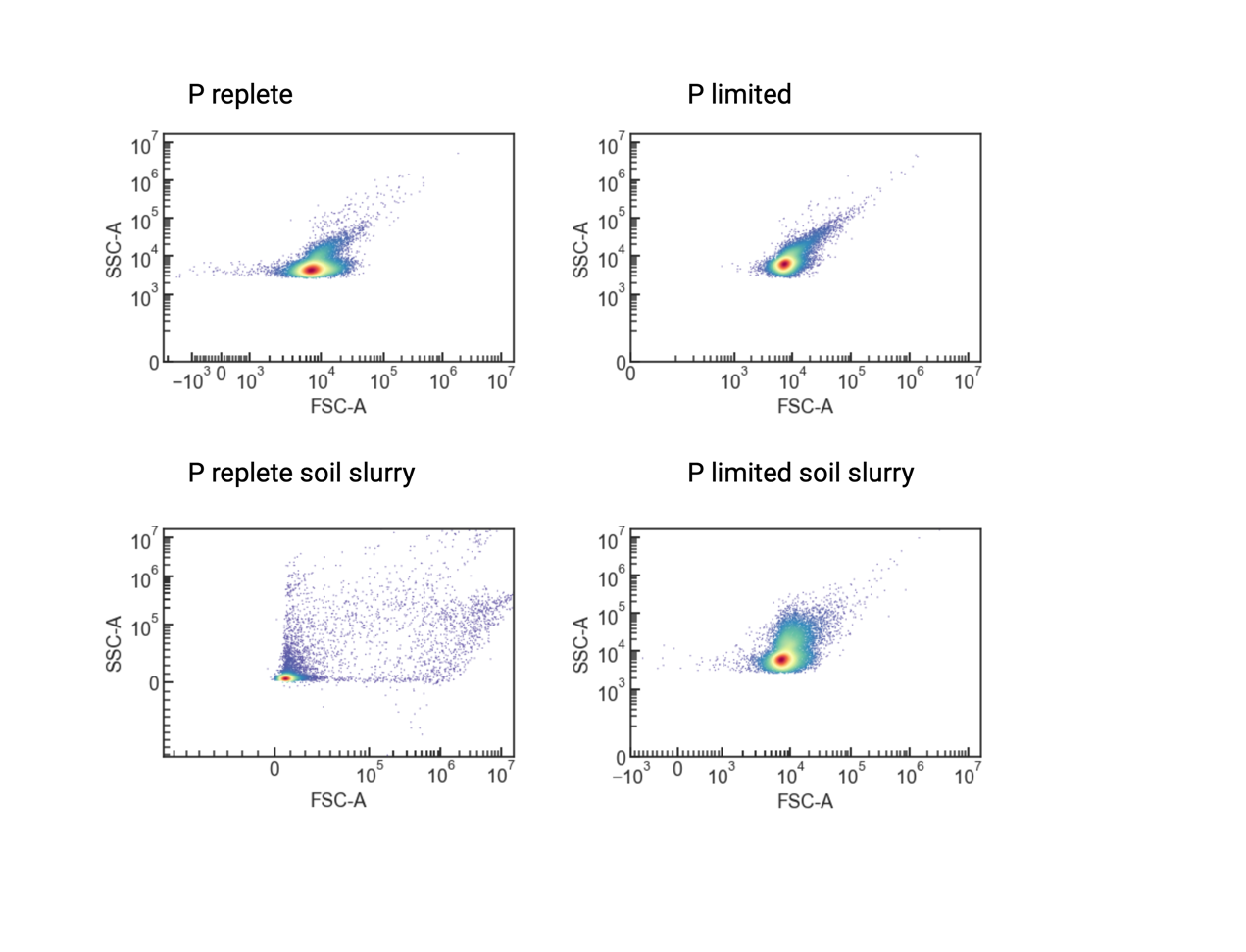

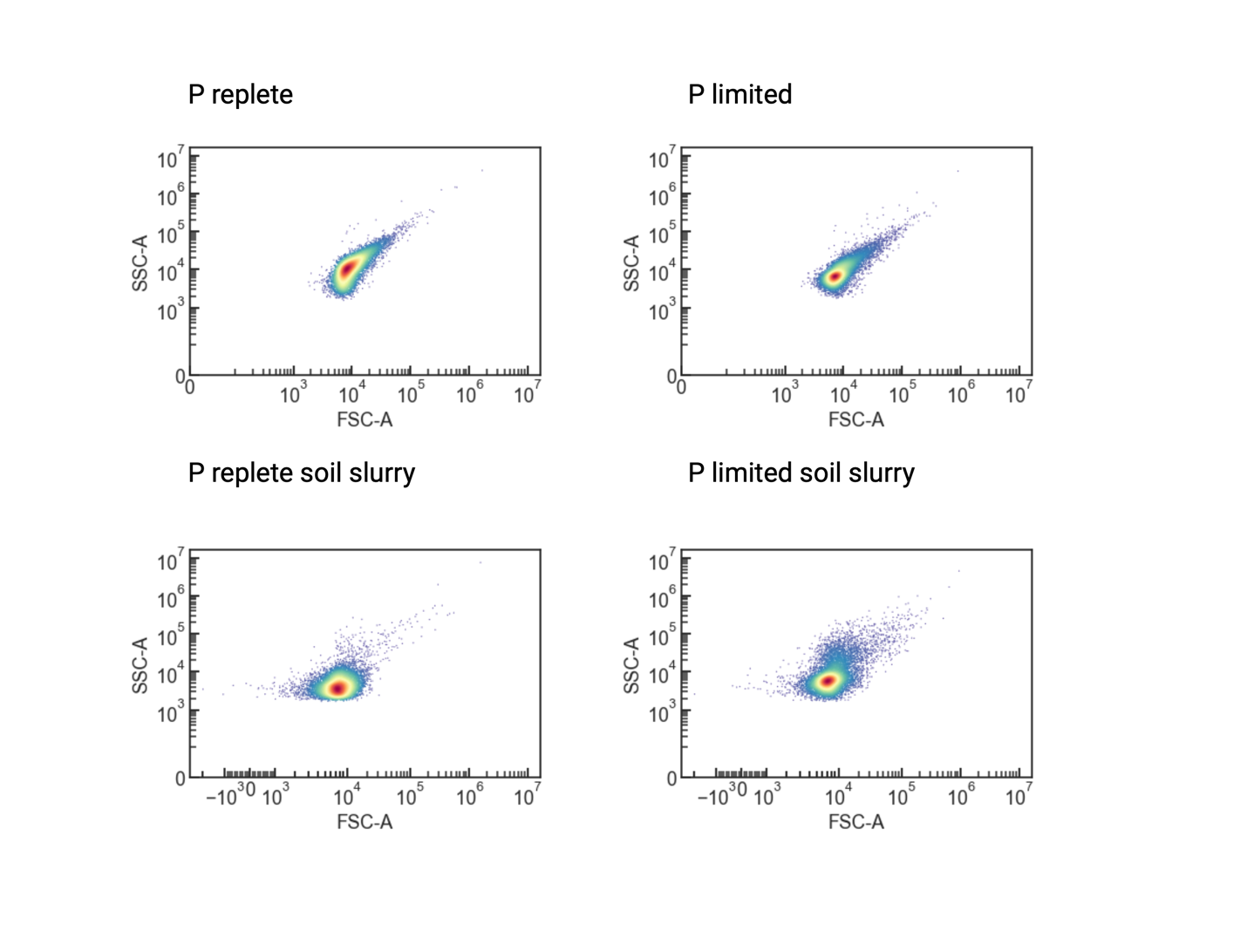

**Figure S5:** Raw flow cytometry data (forward and side scatter) for all three biological replicates.

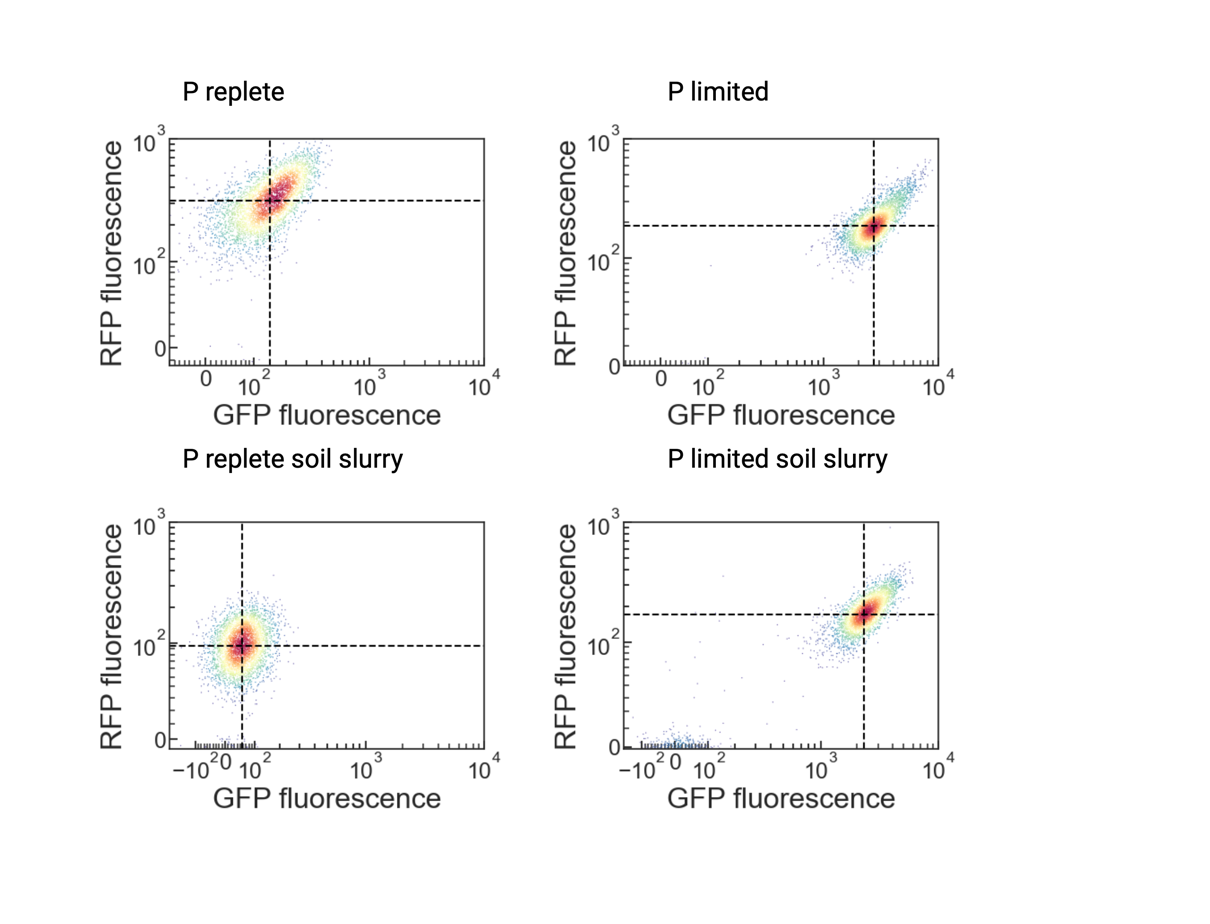

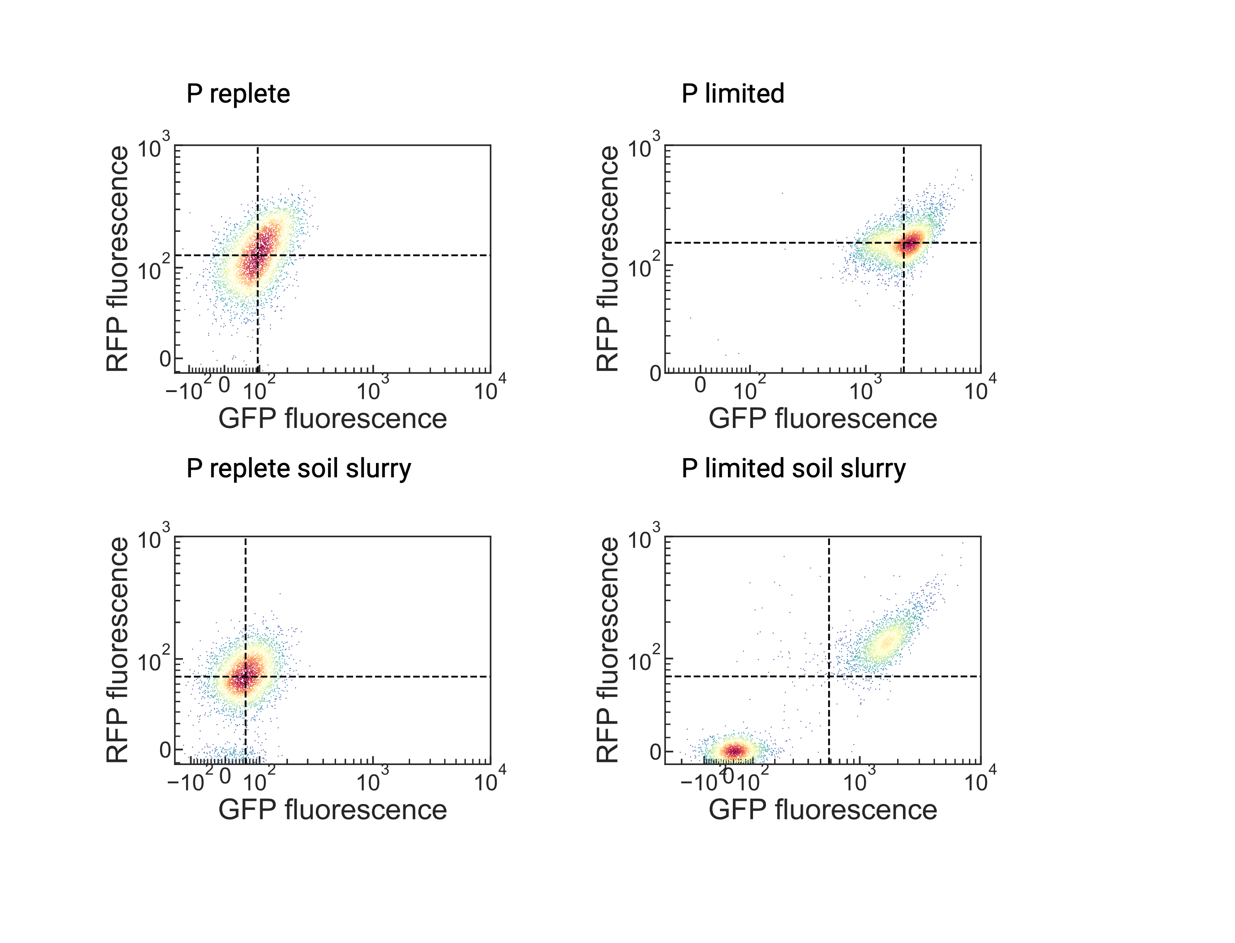

**Figure S6:** Gated flow cytometry data, dashed lines indicating the calculated median fluorescence values.

**References**

[1] R. D. Monds, P. D. Newell, J. A. Schwartzman, and G. A. O’Toole, “Conservation of the pho regulon in *Pseudomonas fluorescens* Pf0-1,” Appl. Environ.
Microbiol., vol. 72, pp. 1910–1924, Mar. 2006.
